# Proteome Complexity Scales with Architecture in Human Neural Models

**DOI:** 10.1101/2025.10.10.681504

**Authors:** Michele Rodrigues Martins, Guilherme Ferraz Alvino, Pedro de Lima Muniz, Christiane Martins de Vasconcellos Silveira, Carolaine da Silva de Araújo, Fábio César Sousa Nogueira, Stevens Rehen, Livia Goto-Silva, Magno Junqueira

**Affiliations:** Department of Biochemistry, Institute of Chemistry, Federal University of Rio de Janeiro, Rio de Janeiro, Brazil; D’Or Institute for Research and Education (IDOR), Rio de Janeiro, Brazil; Department of Genetics, Institute of Biology, Federal University of Rio de Janeiro (UFRJ), Rio de Janeiro, Brazil

**Author notes:** Corresponding authors: Magno Junqueira, Livia Goto-Silva.

**Keywords:** Neuroproteomics, Neural differentiation, Cerebral organoids, Neurospheres, SH-SY5Y cells, Neurodevelopment

## Abstract

Mass-spectrometry–based proteomics enables high-throughput identification and quantification of proteins, providing molecular insight into neural development and cellular organization. Applying this approach to *in vitro* systems of increasing architectural complexity, we compared immortalized monolayer cell lines with two tissue-like neural models. Here, we present a comparative proteomic analysis of three human neural models: SH-SY5Y neuroblastoma cells (2D), neurospheres, and cerebral organoids. Protein profiling revealed a stepwise increase in molecular complexity, with enhanced detection of neural-related proteins linked to axon guidance, synapse formation, and GTPase signaling. This trend was most pronounced in tissue-like models, underscoring their suitability for studying neuronal maturation and circuit assembly. Functional enrichment analyses showed progressive acquisition of neurodevelopmental programs, including synaptic vesicle cycling, presynaptic organization, neurotransmitter regulation, and late-stage gliogenesis, accompanied by increased expression of astrocytic and oligodendrocytic markers. Kinase diversity also increased across models, reaching up to 210 regulatory kinases in organoids, many implicated in neural development and degeneration. Gene set enrichment for neurological pathways mirrored this trend, aligning proteomic complexity with disease relevance. Regional brain mapping further indicated that organoids most closely recapitulate the protein architecture of the human CNS. Together, these findings demonstrate that tissue-like neural models provide richer proteomic landscapes that approximate in vivo brain biology and support the application of integrative proteomics in neuroscience and translational research.

## Introduction

The complexity of the human brain and its central role in defining human cognition and behavior have long been a primary interest in neuroscience. Since *in vivo* studies of the human brain face considerable ethical and technological constraints,^1^ *in vitro* human neural cellular models have emerged as indispensable tools for exploring the central nervous system (CNS), enabling detailed investigations of neurodevelopment, neural function, and disease mechanisms at the molecular and cellular level.^2^

Cell line-derived cultures largely contributed to the study of neural cell biology. However, these models lack the structural and functional complexity of the three-dimensional (3D) architecture of the brain.^3^ Historically, the search to reproduce *in vitro* the structures that resemble the tissue of origin for physiologically relevant discoveries began with Aron Moscona’s neural reaggregates in the 1950s–1960s, progressed with the first self-organized cortical tissues derived from embryonic stem cells reported by Eiraku et al. (2008), and culminated in the cerebral organoids first described by Lancaster et al. (2013), which exhibit spontaneous regionalization of brain-like structures.^4–6^

Brain organoids replicate the tissue organization *in vitro* microenvironment, offering improved platforms for studying tissue organization, neural circuitry, and disease pathophysiology^7^. An alternative spheroid culture termed neurosphere is largely used as it matures faster, starting from neural progenitors and providing 3D interactions between different cell types, including progenitors, astrocytes, and neurons. Comparing these two tissue-like systems permits an assessment of how architectural maturation and prolonged culture shape the neural proteome.

Advancements in proteomics, particularly mass-spectrometry (MS)–based methodologies, have revolutionized our ability to analyze proteins on a large scale.^8^ Contemporary neuroproteomic workflows go beyond simple protein inventories, providing tools to map signaling pathways, post-translational modifications, and neuron-specific proteome dynamics in both health and disease.^9^ MS enables the identification and quantification of thousands of proteins, revealing their roles in various biological processes and diseases. The availability of comprehensive proteomic datasets facilitates cross-comparison, reproducibility, and independent validation, contributing to a deeper understanding of complex systems such as the human brain.^10^

Here, we leveraged and integrated proteomic datasets from three distinct human neural cellular models to investigate how increasing model complexity influences the representation of neural pathways. Specifically, we analyzed the following datasets: neuroblastoma, neurospheres, and brain organoids.

Altogether, all models identified proteins linked to neural mechanisms, but there is a noticeable increase in both the number and diversity of these proteins in tissue-like cultures. This comparison shows that neurospheres expand the range of neural processes, while organoids exhibit even more complex profiles. Organoids encompass a broader spectrum of neurodevelopmental processes, synaptic organization, kinase signaling, and disease-associated pathways. These characteristics underscore the importance of tissue-like models as closer representations of the *in vivo* brain environment and as valuable platforms for studying neurodevelopment and neurological disorders.

## Results and discussion

In this study, we conducted a comprehensive analysis of the proteomic signatures across three human neural models, leading to the identification of numerous proteins and peptides. In the neuroblastoma model, 1,742 proteins and 3,656 peptides were identified, corresponding to a total of 12,944 peptide-spectrum matches (PSMs). The neurospheres model showed a marked increase, with 3,958 proteins, 10,139 peptides, and 47,178 PSMs. The most complex dataset was observed in the Organoids model, which identified 6,249 proteins, 38,584 peptides, and 182,189 PSMs. Comparisons were based on Spectral Abundance Factor (SAF)^11^, which enables robust cross-model integration and is widely used for comparative proteomics; details are in the methods.

### Tissue-like models show a marked increase in axon guidance proteins

The chord diagram (Fig. 1A) revealed extensive overlap between the proteomes of neuroblastoma cells, neurospheres, and organoids. Beyond overlapping with the other models, organoids also contain a significant number of unique proteins (2,615), suggesting advanced neural differentiation and complexity. Neurospheres and neuroblastoma showed 463 and 61 unique proteins, respectively, while 1,950 and 1,479 proteins were shared between the two models. These results highlight the progressive increase in proteomic diversity with model complexity, reinforcing the biological relevance of tissue-like cultures in capturing key neural proteins and cellular diversity.^12,13^

**Figure 1.**
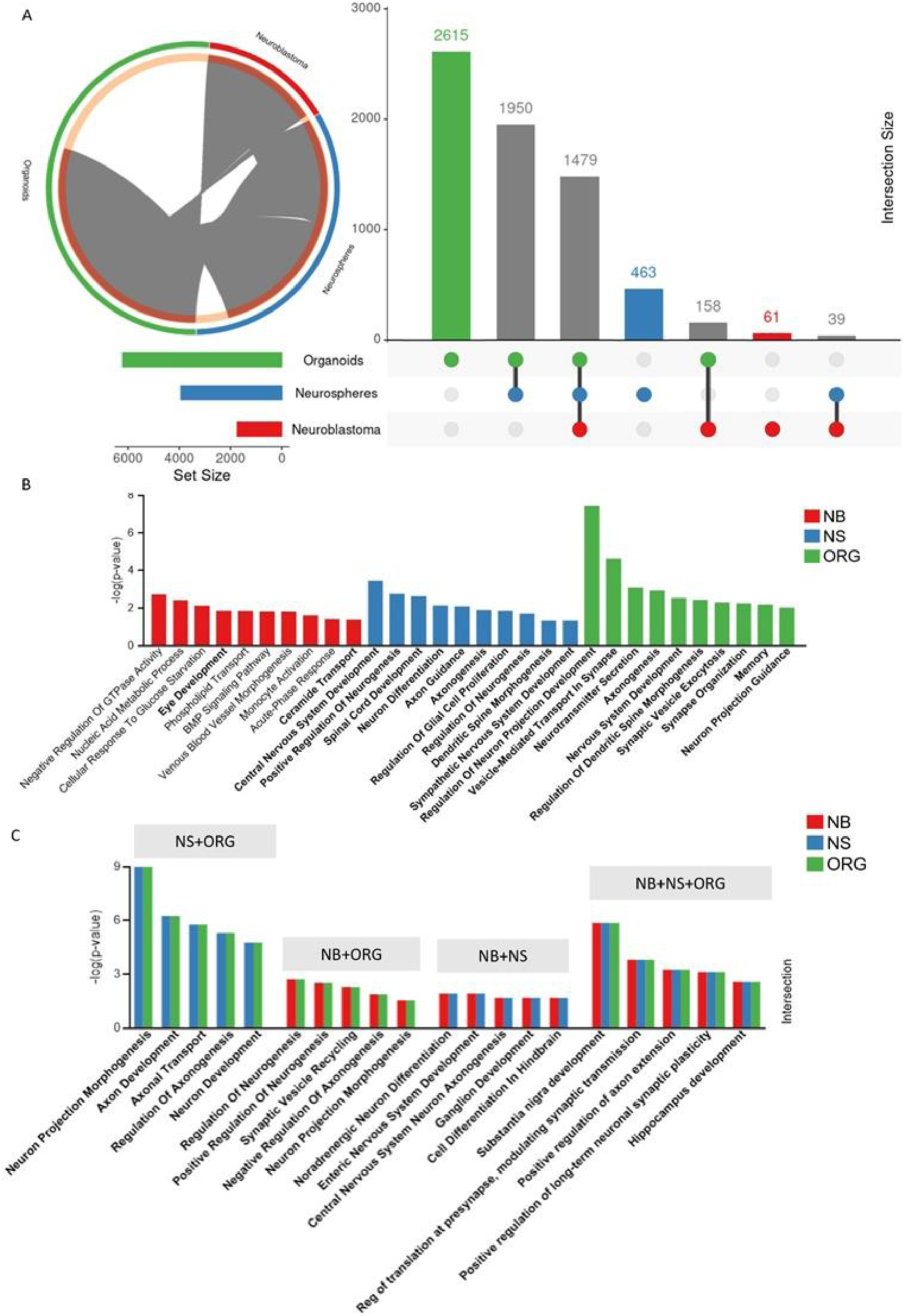
Overview of Identified Proteins. (A) Chord diagram and upset plot showing the intersection of proteins identified in neuroblastoma cells (NB), neurospheres (NS), and cerebral organoids (ORG) with shared and unique proteins in the three models. Highlight for organoids with coverage of a large number of shared and unique proteins. (B) Selected GO enrichment analysis of biological processes identified in the exclusive proteins of each model, with neural-related terms highlighted in bold. (C) Selected GO enrichment of biological processes using protein intersections, also with neural-related terms highlighted in bold.

The GO enrichment analysis demonstrated a clear progression in the complexity of neural processes across the models. Neurospheres showed a broader enrichment of pathways related to neuron projection development, axonogenesis, axon guidance, and synapse organization, suggesting a more structured neuronal network compared to neuroblastoma (Fig. 1B). This is consistent with previous reports that neurospheres exhibit more advanced neural differentiation and network formation, making them suitable for studying neural connectivity and development ^14^. In contrast, neuroblastoma showed minimal enrichment of neuronal pathways in our analysis (Fig. 1B). Among the models, organoids displayed the most complex profile, with significant enrichment in neurogenesis, neuron differentiation, synaptic signaling, and neurotransmitter transport (Fig. 1B).

Neural-related processes were particularly enriched in the shared protein group, highlighting common developmental pathways across different models. Neurospheres and organoids exhibited a broader representation of axon development and guidance, synaptic processes, plasticity, and neurodevelopmental pathways (Fig. 1C). This reinforces the biological relevance of tissue-like cultures in modeling complex molecular mechanisms, including the expression of proteins involved in neuronal maturation and neural circuit formation.^15,16^ These models represent a physiologically relevant platform for examining neurodevelopment, discovering biomarkers, and assessing drug responses in neurological disorders, as evidenced by the enrichment of multiple pathways associated with synaptic organization and axonal remodeling.^17–19^

### Synaptic and membrane trafficking proteins have distinct signatures according to complexity scale

Neuronal communication and brain organization rely on molecular mechanisms that regulate synaptic activity, plasticity, and protein distribution. Understanding how these processes vary across different neural models is essential for evaluating their biological relevance. Figure 2 explores these aspects by comparing neuroblastoma cells, neurospheres, and organoids, providing insights into synaptic representation, protein localization, and neural organization.

**Figure 2.**
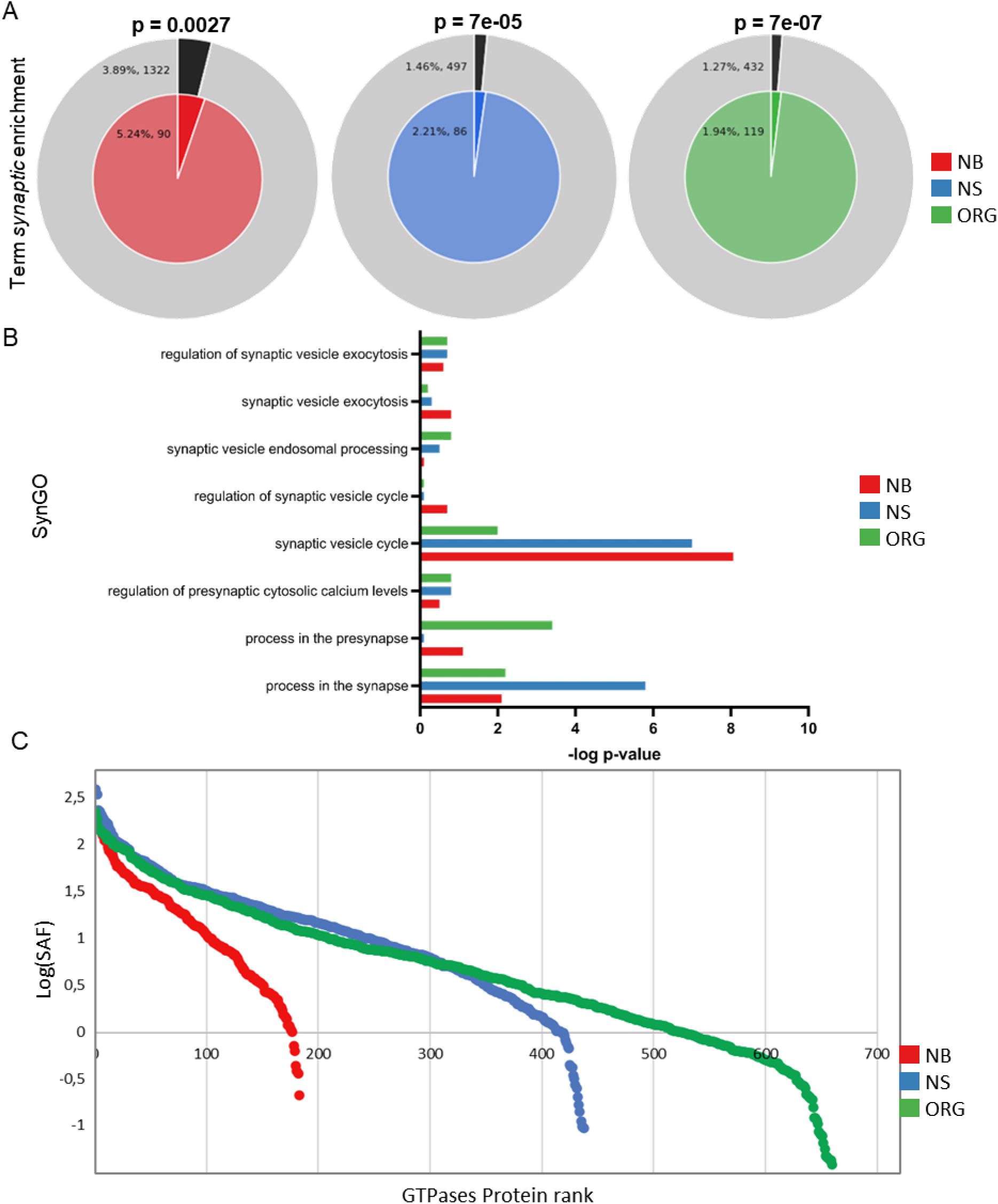
Functional and regional analysis of neuroblastoma in red (NB), neurospheres in blue (NS), and organoids in green (ORG). (A) Enrichment of genes matching the term “synaptic,” with p-values indicating statistical significance. The pie charts display background genes and those present in all models across the GO Biological Processes, KEGG Pathways, and Reactome. (B) Interleaved bars plot representing SynGO enrichment, summarizing the enrichment of synaptic terms across different analyses performed using EnrichR. (C) Ranking of GTPase-related proteins by SAF across neural models.

To assess the representation of synapse-related terms in the datasets, we performed an Association Search for the term “synaptic” in neuroblastoma, neurospheres, and organoids (Fig. 2A). This analysis compares the occurrence of synapse-related genes in each dataset with their genome-wide frequency, applying statistical tests to determine significance. Neuroblastoma exhibited the highest proportion of synapse-related genes (5.24%, 90 genes). This reflects the relative homogeneity of this cell population, where neuronal markers are more represented compared with heterogeneous tissue-like cultures. Consequently, neurospheres (2.21%, 86 genes) and organoids (1.94%, 119 genes) exhibited lower proportions due to the presence of multiple cell types, yet their enrichment was statistically stronger. In all cases, the proportion of synaptic genes was higher than expected relative to the genome-wide background (neuroblastoma: 3.89%; neurospheres: 1.46%; organoids: 1.27%). Statistical analysis confirmed significant enrichment in all datasets (Neuroblastoma: p=0.0027, Neurospheres: p=7e-05, Organoids: p=7e-07). These results indicate that synaptic genes are consistently overrepresented across all models, with a stronger representation in tissue-like cultures. Although neurospheres and organoids contained a smaller percentage of synaptic genes, the enrichment was statistically stronger. This is because, in these more heterogeneous models, the presence of synaptic genes stands out more clearly against what would be expected by chance, resulting in lower p-values.

To investigate synaptic processes, we performed a SynGO-based enrichment analysis via Enrichr (Fig. 2B). SynGO is a knowledge base that classifies synaptic genes and proteins based on experimentally validated evidence.^20^ The analysis focused on shared synaptic genes, with an interleaved bar graph to illustrate the distribution of enriched SynGO terms.

In the neuroblastoma model, we observed an association with the term synaptic vesicle cycle and a stronger signal in synaptic vesicle exocytosis, demonstrating that neuroblastoma cells can express synapse-related proteins. However, this model has important limitations; the high enrichment may be related to the high expression of vesicular genes in a homogeneous tumor lineage, which increases its enrichment even with lower maturity of circuits. In neurospheres, there is an association with terms of regulation of the synaptic vesicle cycle and regulation of levels of presynaptic cytosolic calcium, which indicates the capacity of the model to undergo regulation of synaptic activity, since calcium ions are involved in the regulation of neurotransmitter release during action potentials.^21^ Cortical spheroids present functional neuronal circuits and can also be modulated by pharmacological compounds that affect synaptic transmission.^22^ In organoids, mature synaptic processes such as endosomal processing of synaptic vesicles, presynaptic organization, and regulation of synaptic transmission are more enriched. These findings align with organoid studies showing diverse neuronal cell types and network activity, and suggest that they develop more dynamic synaptic networks ^23,24^

The progressive enrichment of synaptic pathways observed across neuroblastomas, neurospheres, and organoids is closely tied to the function of small GTPases, which are key regulators of synapse formation and maintenance. Synaptic vesicle cycling, presynaptic organization, and calcium regulation all require precise control of actin and microtubule dynamics, processes that are orchestrated by families of GTPases such as Rho, Rab, and Ras. These molecular switches govern vesicular trafficking, endocytosis, and exocytosis at the presynaptic terminal, while also modulating dendritic spine morphology and postsynaptic plasticity. ^25,26^

To explore the distribution of GTPase-related proteins in our neural models, we identified all GTPases using UniProt annotations and ranked them by their SAF in Figure 2D. The analysis revealed differential abundance patterns among neuroblastomas, neurospheres, and organoids, suggesting progressive maturation of cytoskeletal regulation, with increased representation of synaptic GTPases in organoids. GTPases act as signalers that orchestrate neuronal differentiation, cytoskeletal remodeling, and synaptic plasticity. By switching between GDP (inactive) and GTP (active), they coordinate axonal guidance, dendritic morphogenesis, and vesicle trafficking. ^27,28^

Dysregulation of these proteins has also been associated with neurodegenerative diseases. RAB3A, DNM1 and DNM2, essential for endocytosis and mitochondrial dynamics, were more abundant in organoid^29^. Several of the proteins such as RHOA and its effector kinase ROCK2, participate in pathways that drive axonal degeneration and neuronal death in Alzheimer’s and Parkinson’s disease.^30^

Prominent Rho family GTPases (RHOA, RAC1, and CDC42) were detected and are regulators of neuronal growth and connectivity.^31^ They regulate the dynamics of the actin cytoskeleton, promoting the growth and maintenance of dendritic spines and the formation of synaptic contacts.^32^ In addition to Rho-GTPases, we identified multiple members of the septin GTPase family, which also contribute to neuronal differentiation, guiding the formation of axons and dendrites, and stabilizing postsynaptic densities and dendritic spines.^33^ Overall, GTPase profiles show a clear progression across models: neuroblastoma presents a restricted set, dominated by ROCK2 and septins; neurospheres display an intermediate pattern, with moderate levels of RHOA, RAC1 and CDC42; and organoids exhibit the broadest repertoire, including high levels of RHOA, RAC1, CDC42, RAB3A, DNM1, DNM2 and multiple septins, consistent with advanced synaptic organization and increased neuronal connectivity in organoids.

### Pathways, Patterns, and Regional Identities across models show multiple region enrichment in brain organoids

Neurodevelopment is a complex process involving coordinated molecular mechanisms that regulate neuronal differentiation, projection development, and regional patterning. Understanding how these pathways vary across different models provides insight into their ability to recapitulate key aspects of brain organization. Figure 3 explores these features by examining neurodevelopment-related gene representation, enriched biological processes, and tissue-specific expression patterns across neuroblastoma cells, neurospheres, and organoids.

**Figure 3.**
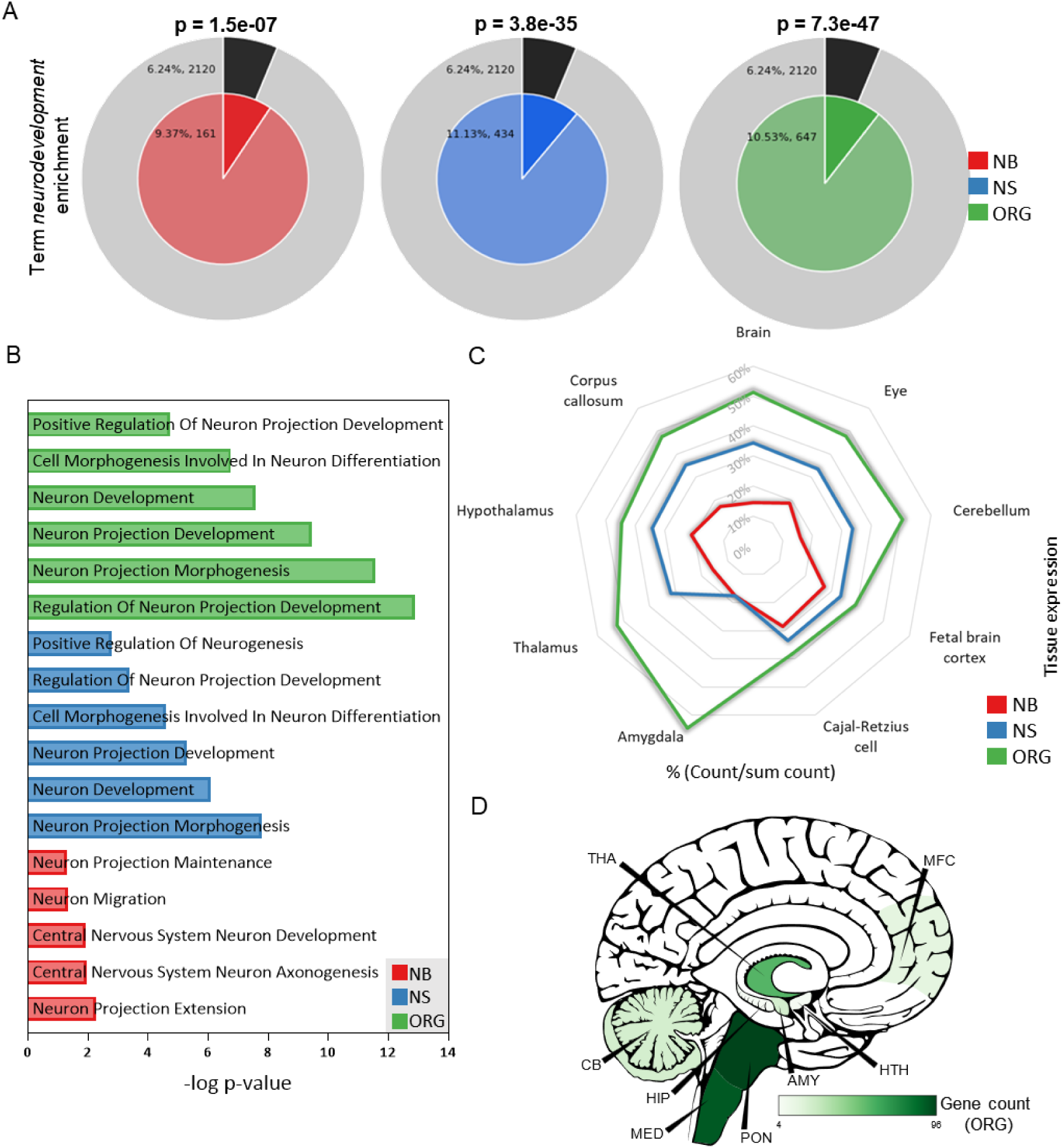
Overview of neurodevelopment-related gene enrichment and tissue distribution in neuroblastoma cells (NB), neurospheres (NS), and cerebral organoids (ORG). (A) Enrichment of genes associated with the term neurogenesis, identified using GO Biological Processes, KEGG Pathway, and Reactome Gene Set (B) GO biological processes related to neurodevelopment that are enriched in the datasets. (C) Distribution of tissue across datasets for each brain region identified in all three datasets (NB, NS, ORG), where the gene count was normalized by the total number of genes mapped to that region across the three datasets, allowing for all direct comparison of relative representation.(D) Brain region mapping of Organoids using CerebroViz, illustrating the areas with the regionally elevated proteins based on Protein Atlas data, where shades of green (organoid model) represent the number of genes associated with each brain region.

To evaluate how neurodevelopment-related genes are distributed across different neural models, we analyzed the occurrence of the term “neurodevelopment” in neuroblastoma, neurospheres, and organoids (Fig. 3A). The proportion of genes associated with this term increased in tissue-like models, with neuroblastoma containing 9.37% (161 genes), neurospheres 11.13% (434 genes), and organoids 10.53% (647 genes). In all cases, these values exceeded the expected genome-wide background (6.24%, 2120 genes). Statistical analysis confirmed that all datasets were significantly enriched, with neuroblastoma (p = 1.5e-07), neurospheres (p = 3.8e-35), and organoids (p = 7.3e-47). The marked enrichment in neurospheres and organoids suggests that tissue-like models provide a broader representation of neurodevelopmental pathways, likely due to their increased cellular complexity and differentiation potential.

Given the enrichment of neurodevelopment-related terms observed in Figure 3A, we next investigated which GO Biological Processes were overrepresented in each dataset (Figure 3B). In neuroblastoma, the enrichment has some critical terms for early stages, such as neuron projection extension and axonogenesis of CNS neurons (–log p = 2.27 and 1.94, respectively), accompanied by terms related to neuronal migration and projection maintenance. Because neuroblastoma arises from neural crest–derived sympathoadrenal precursors,^34^ tumors can fluctuate between adrenergic and mesenchymal/neural crest–like lineage states^35^ and reuse some axon guidance pathways.^36,37^ This helps explain the appearance of these terms in our set. In the Neurosphere analysis, structural remodeling processes emerge, with strongly enriched terms such as neuron projection morphogenesis (–log p = 7.78), neuron development (–log p = 6.1), and cell morphogenesis involved in neuron differentiation (–log p = 4.6), reflecting an increase in complexity marked by the onset of branching and neuronal differentiation in neurospheres.^38,39^ In Organoids, this gradient intensifies. The most significant terms are regulation of neuron projection development (–log p = 12.89), neuron projection morphogenesis (–log p = 11.56), neuron projection development (–log p = 9.45), neuron development (–log p = 7.58), and cell morphogenesis involved in neuron differentiation (–log p = 6.76). This profile suggests not only neurite growth but also tighter control over elongation, guidance, and branching. This shift is consistent with cellular diversity and recent live-imaging data showing morphodynamics during organoid maturation.^24,40^

As the model complexity increases, there is a shift toward the activation of pathways involved in neuronal maturation, structural remodeling, and circuit formation, characteristic of the final stages of neurodevelopment. Similar processes have been described in brain organoids cultured for more than six months, in which the expression of genes linked to axonogenesis, axonal guidance and synaptogenesis progressively increases, concomitant with the appearance of action potentials and mature synapses.^24^ Analogous findings have been observed in cortical spheroids^41^ and in forebrain assembloids that allow functional integration of circuits.^42^ Taken together, the data reveal a clear progression: from the recruitment of basic extension pathways in neuroblastoma, through activation of morphogenesis in neurospheres, to the sophisticated regulation of projections observed in organoids.

Based on the gene and process alterations observed in panels 3A–B, we compared the relative distribution of genes associated with neurodevelopment in ten brain regions common to the three sets (Fig. 3C). For each region, the number of genes was normalized by the sum of the genes from that region in the three models; thus, the percentages reflect the contribution of each model and not the absolute total of genes.

In neuroblastoma, genes annotated with the generic term “brain” total 977 hits (14%), while limbic or commissural signatures remain modest, with the amygdala at 18% and the corpus callosum at 17%. These enrichments likely reflect the overlap of neurodevelopmental genes between the central and peripheral nervous system lineages, rather than their origin in the neural crest and the formation of the peripheral sympathetic nervous system, due to the low specificity of the database annotations. In neurospheres, the profile becomes intermediate: the “brain” tally more than doubles to 2,315 genes (34%), the cortical fraction (cerebrum + cerebellum) likewise reaches 34%, and hippocampal genes emerge at 32%, indicating progressive three-dimensional regionalization. In organoids, a much more diversified spectrum appears: the “brain” contingent rises to 3,458 genes (51%), cortical markers climb to the same 51%, and regions linked to commissural projections, such as the corpus callosum, expand to 48%. This increase corroborates the formation of cortical layers and commissural bundles reported in long-term organoids.^6^ The hippocampal fraction reaches 338 genes (53%), consistent with single-cell data identifying hippocampal-like neurons in six-month organoids^24^. The relative contribution of the amygdala rises to 65%, in line with limbic clusters detected by scRNA-seq in 3–6-month organoids^24^ and with the presence of ventral territories described in prolonged organoid cultures.^6^ Assembloid studies further demonstrate that ventral domains can migrate and establish functional connections with cortical regions,^42,43^ reinforcing the feasibility of modeling limbic circuits in 3D. Finally, recent reviews highlight the application of these systems to the study of sex differences and neuropsychiatric disorders.^44^

Additionally, we conducted a brain-region mapping analysis using cerebroViz (Fig. 3D) to compare organoid proteins with Protein Atlas data, identifying the most highly expressed proteins in each anatomical region of the human brain. By leveraging this specialized database, we achieved accuracy in assessing the extent to which the organoids mimicked the CNS, based on the analysis of the identified proteins and their physiological correlations.

Organoids were selected for brain-region mapping due to their regionalization and cellular diversity. As a result, proteins were mapped onto the neurograph and distributed across multiple brain structures, including the thalamus (THA), medulla (MED), cerebellum (CB), hippocampus (HIP), and amygdala (AMY). Protein signatures distribution across these regions reflects fundamental developmental processes and functional specialization.

In CB, we detected ZIC1 and ZIC2, transcription factors that play a critical role in the development and foliation of cerebellar granule cells.^45^ The HIP region exhibited proteins linked to synaptic plasticity and memory formation. A calcium/calmodulin-dependent protein kinase, CAMK2A, serves as a critical regulator of long-term potentiation and synaptic strength. Furthermore, a calcium-binding protein, SCGN, is vital for signaling pathways and facilitating neurotransmitter release.^45,46^ In THA, proteins involved in the patterning of development are enriched. Members of the HOX family (HOXA5, HOXB7, HOXB4) play a fundamental role in defining regional identity during early stages of neurodevelopment.^47^

Other brain regions were also analyzed, with protein identification in areas such as the white matter (WM), basal ganglia (BG), midbrain (MB), and corpus striatum (CP). The Supplementary table S5 shows the complete analysis of these brain regions, with protein counts representing each region.

### Cellular markers of maturity shift related to complexity

The three models presented, neuroblastoma, neurospheres, and organoids, showed 38, 105, and 137 glia-associated genes, respectively (figure 4C). This rising number expresses the evolution and complexity of the analyzed models. Vimentin (VIM) is a type III intermediate filament that plays a role in migration, proliferation and cell division, and its abundance showed a marked increase from one model to the other (NB < NS < ORG). Such escalation is consistent with VIM’s early activation during differentiation and its later roles in axonal regrowth, myelination and neuro-inflammation.^48,49^

**Figure 4.**
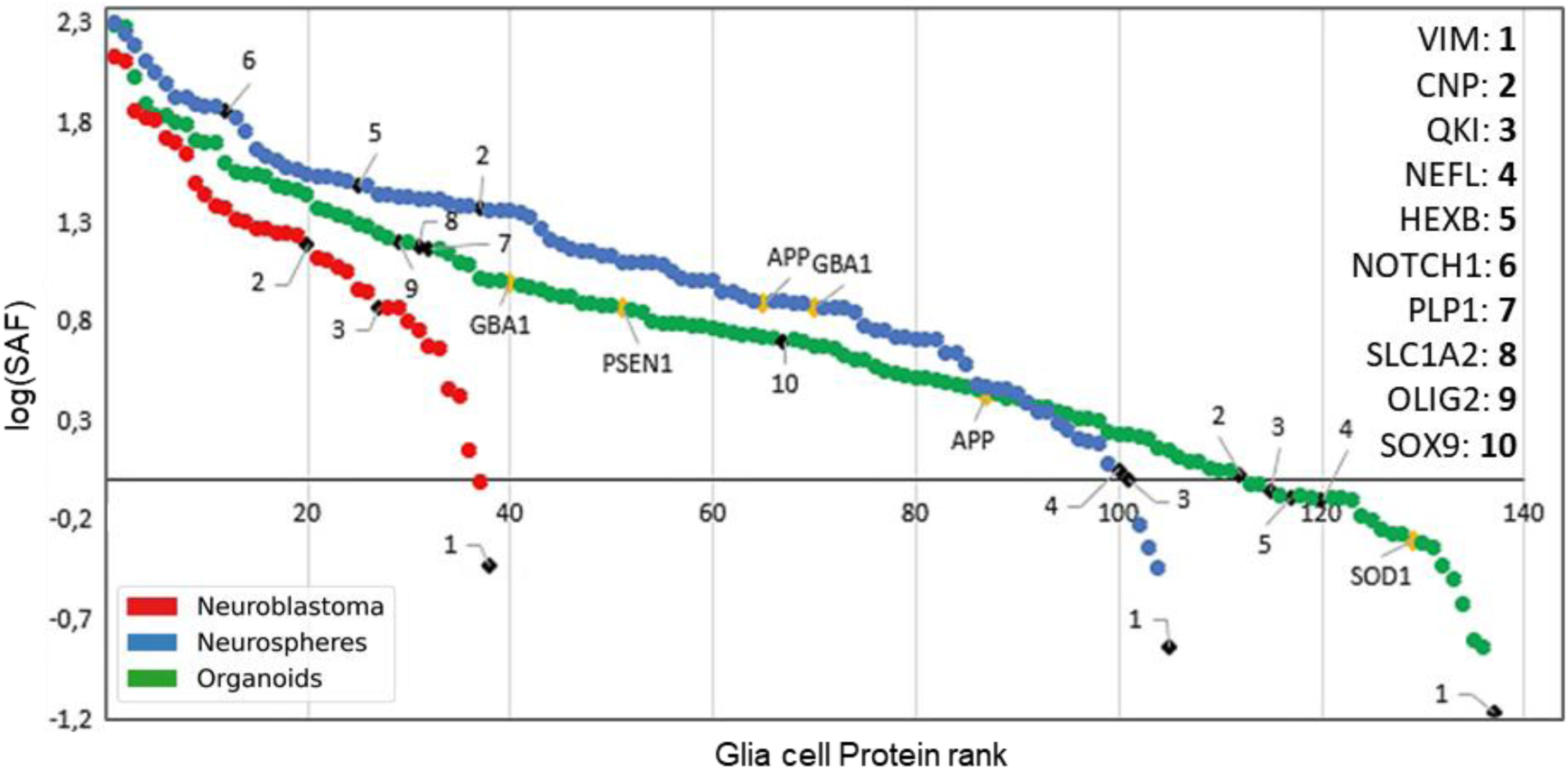
Ranking of glia-associated proteins by SAF across neuroculture models. Black labels denote proteins characteristic of oligodendrocytes (CNP, PLP1, OLIG2) or astrocytes (VIM, QKI, SLC1A2, SOX9, NOTCH1). HEXB, also marked in black, is a lysosomal subunit enriched in microglia in vivo; although its expression suggests immune-lysosomal activity in tissue-like models, it does not confirm the presence of microglia. Yellow labels indicate proteins previously associated with the development of neurodegenerative diseases.

The more complex the neural model, the greater the number of genes reflecting advanced stages of glial development, which explains the detection of SLC1A2 and SOX9 only in the organoid model (Table 1). SOX9 is vital for initiating the neurogenic-to-gliogenic switch in germinal zones and for maintaining neural stem cells.^50,51^ GLT-1, encoded by SLC1A2, rises only post-natally, supporting glutamate clearance in the astrocytic glutamate-glutamine cycle.^52^ The simultaneous detection of SOX9 and the later GLT-1 therefore places organoids at a post-natal-like astrocytic stage that is absent in 2D cultures.

**Table 1.**
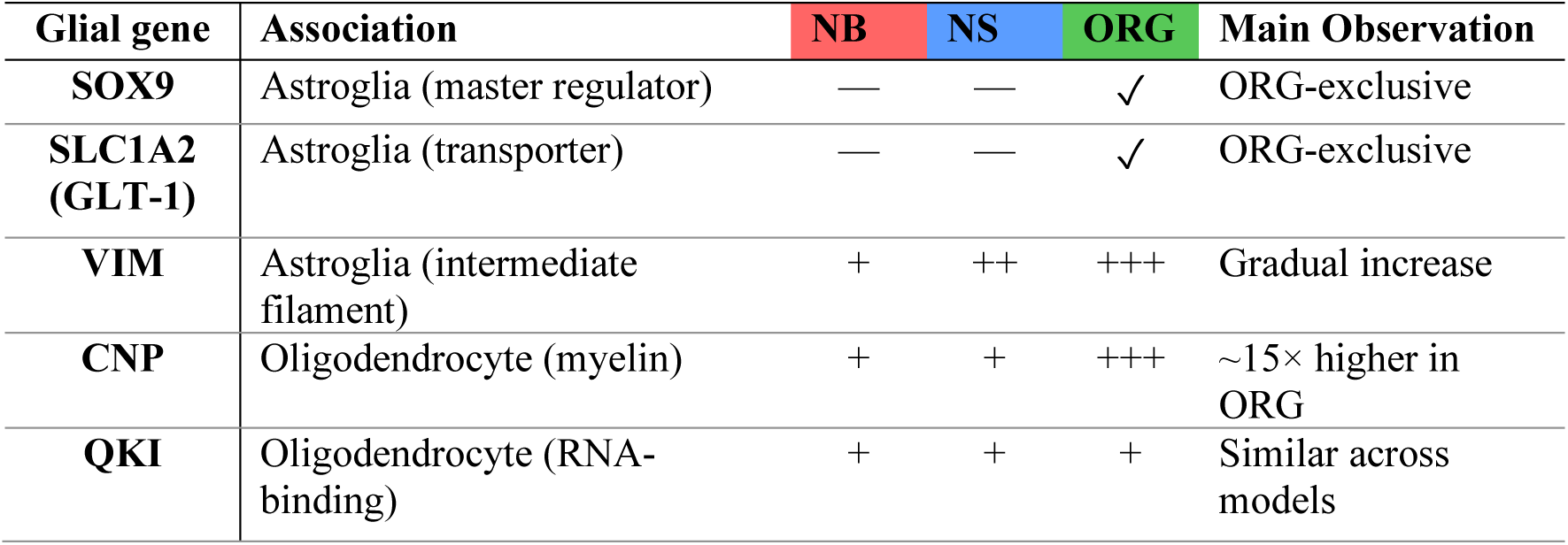
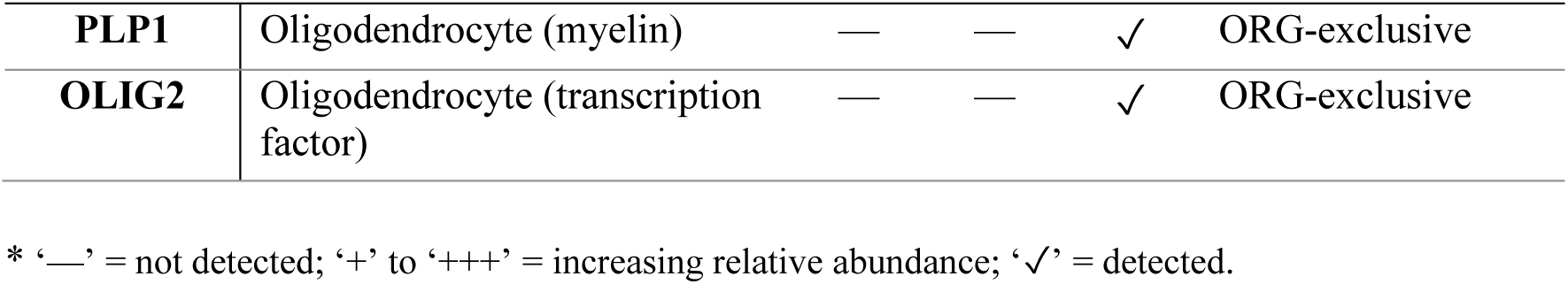
Glial and neurodegeneration markers across the three models.

| Glial gene | Association | NB | NS | ORG | Main Observation |
| --- | --- | --- | --- | --- | --- |
| **SOX9** | Astroglia (master regulator) | — | — | ✓ | ORG-exclusive |
| **SLC1A2 (GLT-1)** | Astroglia (transporter) | — | — | ✓ | ORG-exclusive |
| **VIM** | Astroglia (intermediate filament) | + | ++ | +++ | Gradual increase |
| **CNP** | Oligodendrocyte (myelin) | + | + | +++ | ~15× higher in ORG |
| **QKI** | Oligodendrocyte (RNA-binding) | + | + | + | Similar across models |
| **PLP1** | Oligodendrocyte (myelin) | — | — | ✓ | ORG-exclusive |
| **OLIG2** | Oligodendrocyte (transcription factor) | — | — | ✓ | ORG-exclusive |

\* ‘—’ = not detected; ‘+’ to ‘+++’ = increasing relative abundance; ‘✓’ = detected.

For the oligodendrocyte lineage, CNP and QKI were present in all three models, but the cerebral organoids presented a value approximately 1500% higher for CNP, while PLP1 and OLIG2 appeared only in brain organoids. This pattern mirrors the chronological order of myelin-gene expression, in which CNP precedes PLP, MAG and MOG,^53,54^ underscoring that organoids have entered an early-myelination stage unmatched by neurospheres or neuroblastoma.

### Kinase diversity and ranking evolve with model complexity

Kinase signaling underpins neural development and disease, and its detectable breadth grows as *in vitro* brain models gain cellular and circuit complexity. Notably, many of the neurodevelopmental and neurodegenerative processes are orchestrated by complex signaling networks in which kinases play a central role. Kinases are proteins involved in regulation of neural development, as well as plasticity and cellular homeostasis in the CNS. Dysregulation of this signaling is a common feature of congenital syndromes, neurodegenerative diseases, and brain tumors. In this study, we employed approaches to the main kinase-mediated signaling pathways.

To capture this progression, we first compared the kinases identified across Neuroblastoma, Neurospheres, and Organoids, showing that organoids harbor by far the largest and most diverse repertoire. The Euler diagram (Figure 5A) compares the set of kinases found concomitantly or exclusively in the models, indicating that signaling diversity expands with model maturation. Organoids predominate in number with 210 kinases, 96 of which are unique.

**Figure 5.**
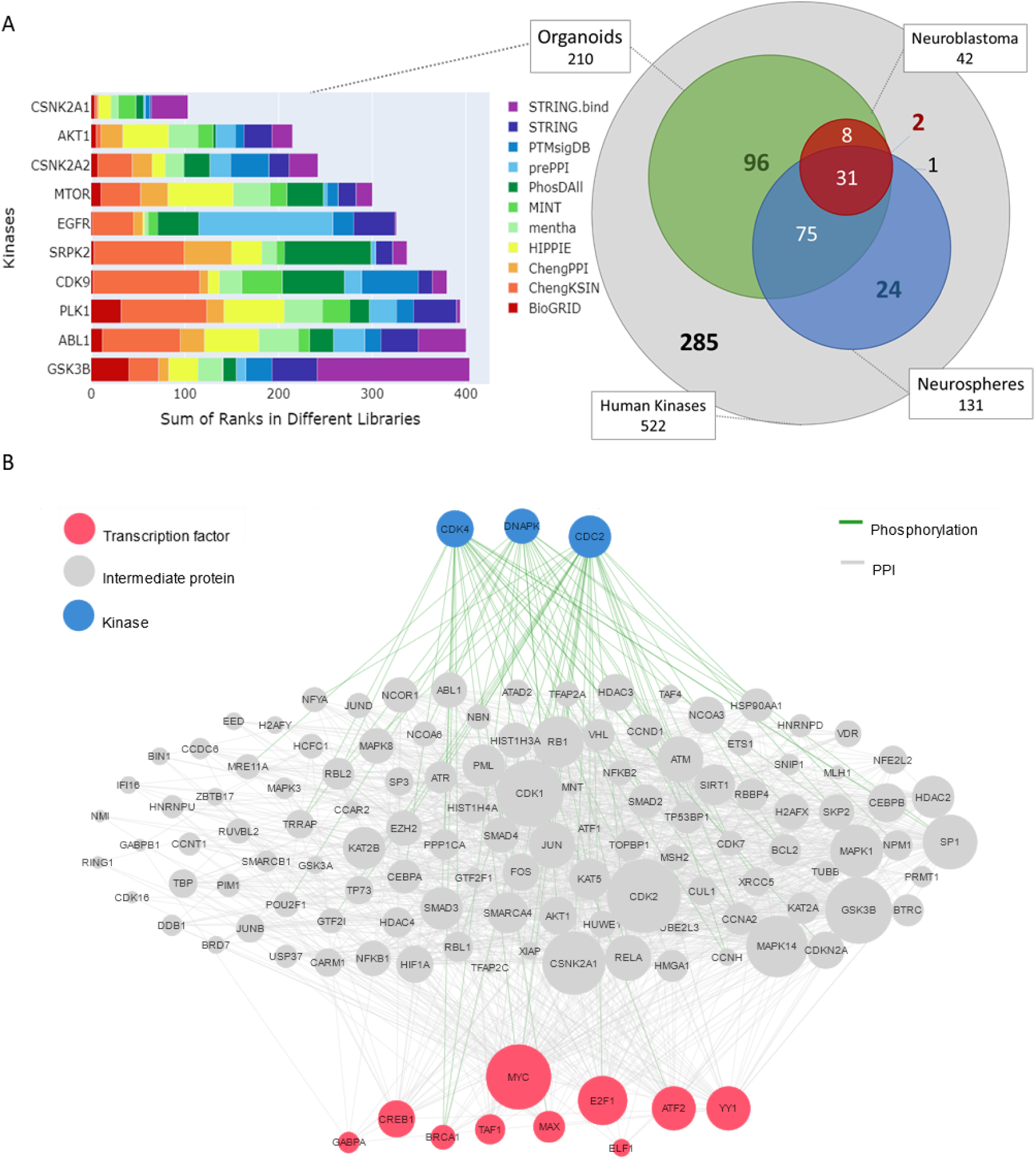
(A) Bar chart (left) showing the average kinase rank in Organoids generated by KEA3, highlighting the most relevant predicted enzymes across libraries, and proportional Venn diagram (right) depicting the overlap of identified kinases in models obtained from the human kinome. (B) Signaling network produced by the X2K algorithm connecting transcription factors (red nodes) to kinases (blue nodes) for the organoids dataset. Gray nodes represent proteins intermediate to the interaction. White edges indicate protein-protein interactions, and green edges represent kinase phosphorylation events.

**Figure 6.**
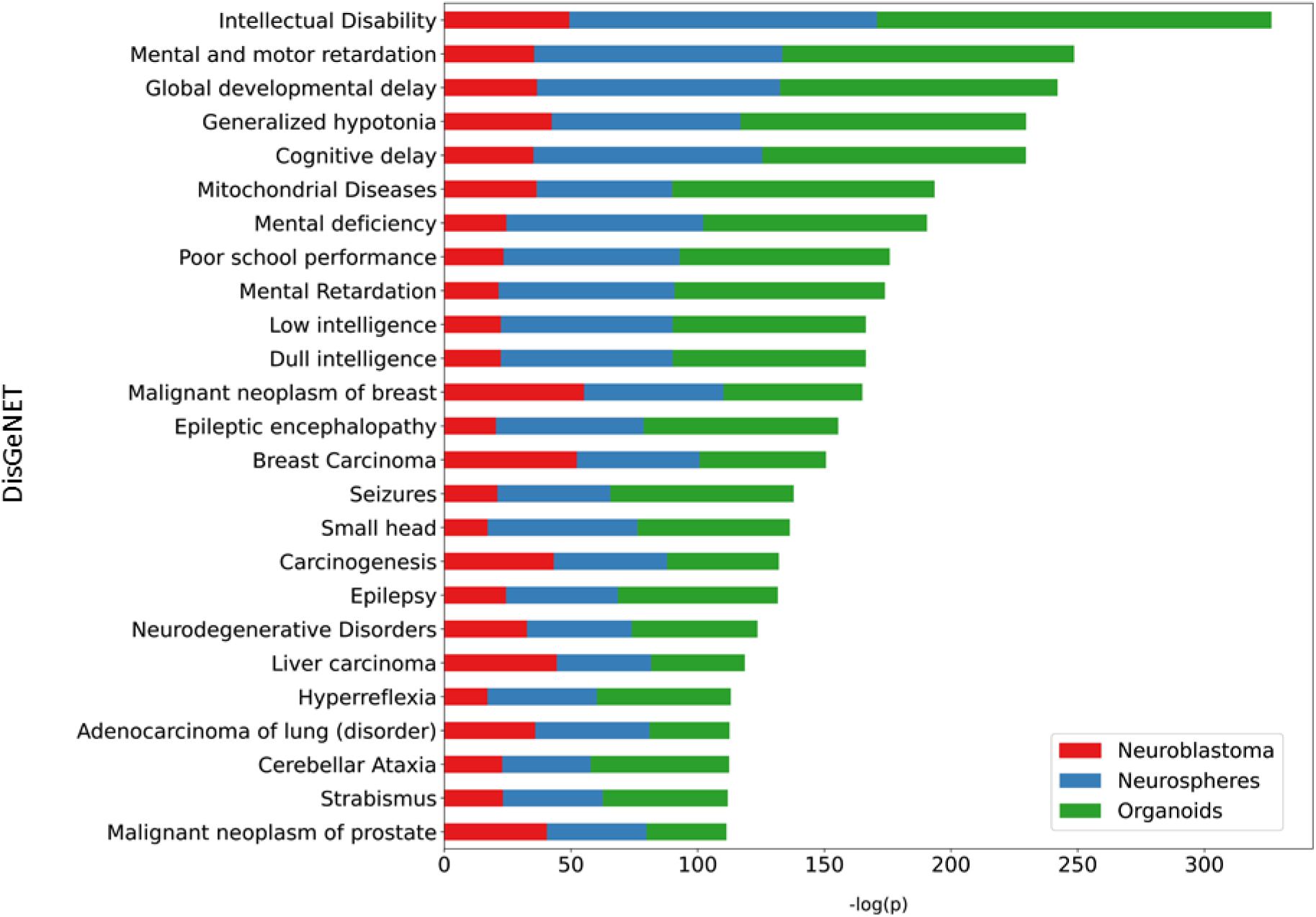
Stacked bar plot for the top 25 consensus terms. Consensus scores are computed by taking the sum of each enrichment analysis of 3 gene models (-log p-value) using the DisGeNET library in EnrichR.

We then focused on organoids for deeper regulatory inference. Using KEA3 MeanRank,^55^ which integrates 11 kinase-substrate libraries, we ranked kinases. The bar graph shows the top five, CSNK2A1, AKT1, MTOR, EGFR and GSK3B, known for their roles in signaling, growth and pathogenesis of neural diseases. mTOR is a central regulator of cell growth and survival. In the CNS, it regulates neuronal survival and differentiation, dendritic arborization, axonal growth, synaptogenesis and synaptic plasticity.^56^ The PI3K/AKT/mTOR axis regulates cell proliferation, survival, metabolism, and differentiation, being essential for neural development and maintenance of pluripotency in stem cells.^57^ AKT target, GSK3B, is involved in the regulation of axonal growth and neuronal plasticity.^58^ The EGFR receptor plays roles in the regulation of neuronal differentiation, neuroglial function, regeneration after injury, and neurodegenerative diseases.^59^

Figure 5B shows a network analysis generated by the X2K algorithm^60^ of the organoids, connecting transcription factors (red nodes) to kinases (blue nodes) and intermediate proteins (gray nodes). Green edges indicate phosphorylation events, and white ones represent protein-protein interactions. At the top of the network are kinases such as CDK4, DNAPK (PRKDC), and CDC2 (CDK1), each connected to numerous intermediate proteins. CDK4 and CDC2 control cell cycle progression of neural progenitors, while DNAPK integrates the DNA damage response with chromatin remodeling.^61^ The gray nodes comprise chromatin remodeling proteins, such as ATR, RB1, HDACs, and SWI/SNF components, suggesting that phosphorylation of these molecules directly influences transcriptional activity.^62^ At the base of the network, the transcription factors MYC, E2F1, BRCA, and CREB stand out. The first regulates growth and metabolism of neural progenitors,^63^ the second coordinates entry into the S phase of the cell cycle^64^, the third is essential for the development and protection of the nervous system^65^, and the last modulates genes linked to synaptic plasticity.^66^ Alterations in the activity of transcription factors is a common feature in neurodegenerative diseases, contributing to neuroinflammatory conditions and oxidative stress.^67,68^

### Molecular profiles associated with neurological diseases and disorders

By integrating proteomic analysis of neural models, we were able to map molecular profiles associated with neurological diseases and disorders. In the DisGeNET disease term graph (Fig. 4A), neurological and developmental disorders are particularly prominent among the enriched categories.

The top bar, corresponding to “Intellectual Disability”, exhibits the highest consensus score, highlighting a strong enrichment of genes associated with this condition across the analyzed models. Other neurodevelopmental and cognitive terms, such as “Mental and Motor Retardation”, “Global Developmental Delay”, and “Cognitive Delay”, also rank among the most significantly enriched terms. In neuroblastoma, terms related to “Malignant Breast Neoplasm” and “Malignant Prostate Carcinoma” are observed, reflecting the cancerous origin of this cell line. However, neuroblastoma also contributes, to a lesser extent, to several neurological terms, indicating its partial relevance to neural and developmental pathways. Both organoids and neurospheres display substantial enrichment for neurological disorders. However, organoids show more pronounced representation in terms related to neurodevelopmental conditions compared with neurospheres, particularly for more specific terms. This pattern suggests that tissue-like neural models better recapitulate the molecular landscape associated with complex neurodevelopmental and neurodegenerative disorders.

Neurodegeneration-linked markers followed the same hierarchy. APP and GBA1 appeared in neurospheres, whereas organoids additionally expressed PSEN1 and SOD1. APP misprocessing and lysosomal GBA1 deficits connect gliogenesis to Alzheimer’s and Parkinson’s pathways.^69,70^ PSEN1-bearing microglia exacerbate Aβ pathology,^71^ while mutant SOD1 in microglia accelerates ALS progression.^72^ Taken together, organoids uniquely capture features of late-stage gliogenesis alongside the early molecular hallmarks of neurodegeneration, whereas neurospheres represent an intermediate stage of neural development, and neuroblastomas remain a restricted 2D baseline.

Kinase and Transcription factor analysis highlighted the intersection of disease-related pathways. Four kinases (AKT1, MTOR, GSK3B, and EGFR) cross-reference intellectual disability and multiple neurodegenerative disorders, linking organoids to signaling regulators implicated in Alzheimer’s, Parkinson’s, and ALS.^59,73,74^ Neurodegenerative Disorders term converge with transcriptional regulators enriched in our dataset. TAF1 variants have been to linked X-linked syndromes encompassing intellectual disability, developmental delay, dysmorphic features, and neurological manifestations.^75^ Loss or inactivation of CREB1 in the brain leads to apoptosis of postmitotic neurons and progressive neurodegeneration in regions such as the hippocampus and striatum, with phenotypes similar to those observed in diseases such as Huntington’s and Parkinson’s.^76,77^ It signaling contributes to neuronal death and synaptic deficits in Alzheimer’s.^78^ MYC is involved in neurodegenerative diseases and neuronal repair.^79^ Finally, YY1 is associated with several neurological diseases, with developmental abnormalities and anatomical malformations in the CNS.^80^

## Conclusions

This study demonstrates a increase in proteome complexity along the model continuum from SH-SY5Y neuroblastoma cells to neurospheres and brain organoids. As models gain cellular and architectural sophistication, we observe broader coverage of neurodevelopmental pathways, clearer evidence of glial maturation (e.g., SOX9, SLC1A2, PLP1/OLIG2), and richer synaptic and regional signatures. Kinase inference further expands with model maturity: organoids encompass 210 kinases (96 unique), and KEA3/X2K prioritization converges on CSNK2A1, AKT1, MTOR, EGFR, and GSK3B, regulators central to neuronal survival, neurite outgrowth, synaptogenesis, and plasticity. Notably, these drivers intersect with disease terms spanning intellectual disability, Alzheimer’s disease, Parkinson’s disease, and ALS, reinforcing the translational relevance of organoids as the most informative platform among those tested for target discovery, mechanism-of-action studies, and preclinical screening.

Some limitations should be acknowledged. The integration of independently generated datasets and the use of SAF as a normalization strategy impose inherent constraints on quantitative interpretation. Moreover, the comparison between 2D and 3D models involves distinct cellular origins (neuroblastoma vs. iPSC-derived), which may contribute to some of the observed differences. Finally, our analysis is restricted to the total proteome, focusing primarily on qualitative comparisons across models. These caveats, however, do not diminish the consistency of the observed trends, nor the relevance of the biological insights uncovered, which contribute important advances to the scientific understanding of neural development and model complexity.

Beyond its conceptual value, this dataset serves as a resource for the community. It provides proteomic baselines for neuroblastoma, neurospheres, and organoids; identifies transcription factors, kinases, and intermediate modules to guide pathway-focused hypotheses; and offers metrics for regional, synaptic, and glial coverage that support model evaluation. These elements establish premises for advancing future studies, from prioritizing kinase inhibitors and assessing differentiation to selecting complementary readouts and mapping candidate biomarkers to disease-relevant pathways. Collectively, the dataset is reusable, comparable across laboratories, and directly informative for experimental planning and validation.

## Methods

### Neural Model Generation and Sample Preparation

Three neural models with increasing complexity were analyzed: SH-SY5Y neuroblastoma cells, neurospheres, and cerebral organoids.

**Neuroblastoma** was a dataset previously published in Murillo et al. (2017).^81^ Briefly, two cell populations were prepared for proteomic analysis. The undifferentiated group was harvested during the exponential growth phase under standard culture conditions. To obtain the differentiated group, cells were treated for 15 days with a sequential regimen of retinoic acid (RA) followed by brain-derived neurotrophic factor (BDNF) and other supplements essential to induce a mature neuronal phenotype. Subsequently, cells from both conditions were collected, washed, lysed, and prepared for mass spectrometry analysis according to the published.

**Neurospheres** were a dataset previously published by Goto-Silva et al. (2021)^14^. Generated from human neural stem cells (GM23279A, Coriell Institute) were seeded in ultra-low attachment plates and maintained for 3 or 10 days in differentiation medium to allow for neurosphere formation. Samples were then harvested, washed, and processed for protein extraction and proteomic analysis according to the published protocol.

**Cerebral organoids** All proteomic analyses and datasets for organoids presented are unpublished. Were generated from human iPSCs (GM23279A, Coriell Institute) following the unguided protocol described and detailed in Goto-Silva et al. (2019).^82^ Briefly, iPSCs were dissociated into single cells and aggregated in 96-well ultra-low attachment plates with ROCK inhibitor for embryoid body (EB) formation, followed by neural induction and Matrigel embedding. Organoids were maintained on an orbital shaker with weekly medium changes, and samples were collected at 35, 45, 60, and 75 days of differentiation. The four organoid stages after digestion were labeled with iTRAQ-4plex and analyzed together. Data are available via ProteomeXchange with identifier PXD065667.

### Protein Extraction, Digestion, and iTRAQ Labeling

For all models, samples were lysed in buffer (7 M urea, 2 M thiourea, 50 mM HEPES pH 8.0, 75 mM NaCl, 1 mM EDTA, protease inhibitors), sonicated, and centrifuged to remove debris. Total protein was quantified using the Qubit Protein Assay (Thermo Fisher). 100 μg of protein per sample was reduced (10 mM Tris(2-carboxyethyl) phosphine, 30°C, 1 h), alkylated (40 mM Iodoacetamide, room temperature, dark, 30 min), diluted with 50 mM Triethylammonium Bicarbonate, and digested with trypsin (1:50, 35°C, overnight). Peptides were desalted on C18 microcolumns and labeled with iTRAQ 4-plex reagents (AB Sciex/Merck) as per manufacturer’s instructions. Labeled peptide mixtures were fractionated by hydrophilic interaction liquid chromatography (HILIC) on a TSKGel Amide-80 column, and 26 fractions were collected and pooled according to chromatographic intensity. Samples were dried, resuspended in 0.1% formic acid.

### Mass Spectrometry Analysis

Peptide fractions were analyzed by nanoLC-MS/MS on an LTQ Orbitrap Velos or a Q-Exactive Plus mass spectrometer (Thermo Scientific), both operated in data-dependent acquisition (DDA) mode. Peptides were first loaded onto a trap column (ReprosilPur C18, 2 cm × 150 μm i.d., 5 μm, flow rate 5 μL/min) and subsequently separated on a homemade nano column (ReprosilPur C18, 30 cm × 75 μm i.d., 1.7 μm, flow rate 300 nL/min) using a linear gradient of 5–40% acetonitrile/0.1% formic acid over 105 min, followed by a rapid increase to 95% B. Spectra were acquired in positive ion mode with a spray voltage of 2.5 kV and a capillary temperature of 200 °C. Full MS scans were acquired at 70,000 resolution (m/z 200) in the range of 375–1,800 m/z, and the top 10 most intense ions were fragmented by higher-energy collisional dissociation (HCD, NCE 30) with MS/MS scans recorded at 17,000 resolution. Dynamic exclusion was set to 45 s.

### Data Processing and quantification

Raw data files were processed in Proteome Discoverer 2.1, with peptide/protein identification against the UniProt human database. Strict and relaxed criteria were applied for FDR (<0.01% and < 0.05%, respectively) and PSM validation. For comparative analyses across models and datasets, we did not rely on the previously reported iTRAQ quantifications.^14,81^ To enable comparisons across independently generated datasets with different preparation dates, enrichment configurations, and the presence of isobaric labeling, we applied a spectral-count–based normalization independent of reporter ions. For each protein, abundance was estimated as the number of peptide-spectrum matches (PSMs) divided by its molecular weight, analogous to the Spectral Abundance Factor (SAF/NSAF) approach that corrects for protein size bias and provides a reproducible proxy for relative abundance.^11,83^ This strategy is widely used in label-free proteomics and is conceptually related to emPAI.^84^ Enrichment analyses for neural-related terms were performed independently for each dataset, and significance was evaluated using the hypergeometric distribution. Together, this normalization and enrichment framework allowed harmonization of two previously published datasets with one newly generated dataset, minimizing batch- and label-specific artifacts.^85^

### Bioinformatic and Functional Enrichment Analysis

The final protein lists for each dataset were analyzed using the R language, Excel, and GraphPad Prism software. Functional annotation and pathway enrichment from Gene Ontology (GO), Reactome, and KEGG pathway were performed using DAVID, Metascape, and EnrichR. Synaptic and brain region analyses were performed using SynGO, cerebroViz, and the Protein Atlas database, focusing on neurodevelopmental and disease-relevant pathways. All identified proteins are listed in Supplementary Table S1, complete enrichment results are provided in Supplementary Tables S2–S4, and brain region mapping is shown in Supplementary Table S5.

## Author Contributions

MM, LG-S performed the experiments. MM, PM, GA, and CS performed the data analysis. MM, PM, and GA produced figures. MM wrote the manuscript. MM, GA, PM, CS, CA, LG-S, SR, and MJ contributed to and validated the manuscript. MJ and SR provided resources and acquired funding. MJ coordinated this work. All authors approved the manuscript.

## Acknowledgements

This project was supported by Fundação Carlos Chagas Filho de Amparo à Pesquisa do Estado do Rio de Janeiro (Projec FAPERJ) and Conselho Nacional de Desenvolvimento Científico e Tecnológico (CNPq).

The author(s) used ChatGPT (OpenAI) for sentence review of the writing. The author(s) reviewed and edited the content as necessary and assume full responsibility for the content of the publication.

